# The Aging Microenvironment as a Determinant of Immune Exclusion and Metastatic Fate in Pancreatic Cancer

**DOI:** 10.1101/2025.05.08.652966

**Authors:** Priyanka Gupta, Rabi Murad, Lily Ling, Yijuan Zhang, Karen Duong-Polk, Swetha Maganti, Cheska Galapate, Cosimo Commisso

## Abstract

Aging is a critical yet understudied determinant in pancreatic ductal adenocarcinoma (PDAC). Despite a strong epidemiological association with age, conventional PDAC preclinical models fail to capture the histopathological and stromal complexities that emerge in older organisms. Using an age-relevant syngeneic orthotopic model, we demonstrate that organismal aging accelerates PDAC progression and metastasis. Through transcriptomic profiling, we identify a conserved extracellular matrix gene signature enriched in cancer-associated fibroblasts (CAFs) from aged tumors, consistent with an augmented fibrotic landscape that supports immunosuppression, metastatic tropism, and poor prognosis. To directly test the functional impact of stromal aging, we employed heterochronic co-implantation models, revealing that revitalizing the aged tumor stroma with young CAFs restores immune infiltration and attenuates metastasis in older hosts. Conversely, aged CAFs, while immunosuppressive, fail to enhance metastasis in young hosts, suggesting that a youthful microenvironment exerts dominant regulatory control over disease progression. These findings demonstrate that stromal age is a critical modulator of both immune exclusion and metastatic behavior in PDAC. Importantly, our work establishes a new conceptual framework for understanding how aging shapes the tumor microenvironment in PDAC and opens a fertile avenue of investigation into age-specific stromal regulation. Moreover, this work raises compelling questions about the underlying molecular mechanisms—questions now accessible through our models—and lays the foundation for future efforts to therapeutically target stromal aging in PDAC.

**Statement of Significance:** Our study links aging, stromal remodeling, and PDAC aggressiveness, highlighting how age-dependent stromal changes drive progression and suggesting that rejuvenating the aged microenvironment may improve outcomes in older patients.

## Introduction

Aging is a major risk factor for cancer, yet its impact on disease progression remains underexplored in many malignancies, including pancreatic ductal adenocarcinoma (PDAC). The increasing rates of metastatic PDAC are closely linked to advancing age and poorer outcomes in older adults. Population-based data shows that the median age at diagnosis for PDAC is 71 years, with approximately 65% of cases diagnosed in individuals over the age of 65 (1–3). Despite the clear epidemiological link between advancing age and PDAC incidence, the underlying biological mechanisms that render older individuals more susceptible to disease remain largely elusive—a critical knowledge gap with direct clinical consequences, given that the majority of patients are diagnosed late in life. This persistent void in our understanding likely stems from a fundamental disconnect between clinical reality and the experimental paradigms used to study PDAC, which overwhelmingly rely on young models that fail to capture the complex biological context of aging. For instance, most preclinical research models for PDAC typically rely on young mice aged 2 to 6 months, which is roughly equivalent to 20- to 30-year-old humans thereby failing to recapitulate the molecular and pathological features of PDAC in the elderly. Consequently, effective treatment strategies for this growing patient population remain inadequately defined, undermining efforts to improve outcomes in older adults with PDAC (4–6). Incorporating age as a biological variable in preclinical models holds immense potential to illuminate how the pathological complexity of PDAC—and its trajectories of progression and metastasis—manifests uniquely in older organisms, ultimately guiding the development of more precise and effective interventions for this underserved population.

The stroma in PDAC plays a central role in shaping nearly every aspect of tumor biology, from supporting tumor cell growth and survival to influencing immune evasion and therapeutic resistance. Far from being a passive scaffold, the stromal compartment actively communicates with cancer cells through biochemical and mechanical cues, orchestrating tumor progression and metastasis. In PDAC, the tumor microenvironment consists of a diverse array of cell types— including fibroblasts, immune cells, endothelial cells, and pericytes—embedded within a dense and highly remodeled extracellular matrix (ECM). Among these, fibroblasts are particularly abundant and undergo activation within the tumor microenvironment to become cancer-associated fibroblasts (CAFs), which are key orchestrators of tumor progression. CAFs contribute to a myriad of aspects of tumor biology: they remodel the extracellular matrix to create a supportive niche for tumor growth, secrete growth factors and cytokines that promote cancer cell proliferation and survival, and modulate immune responses to create an immunosuppressive environment. Additionally, CAFs facilitate angiogenesis and generate mechanical forces that promote tumor invasion and metastasis. Systemically, normal fibroblasts are especially susceptible to age-related changes with well-documented functional declines and phenotypic shifts that influence tissue homeostasis (7–14). Notably, normal fibroblasts isolated from aged, non-malignant human pancreas enhance the proliferative capacity of PDAC cells compared to fibroblasts from younger individuals, an effect attributed to differences in their secretomes (15). Age-related changes in normal fibroblasts might alter tumor dynamics in complex and context-dependent ways since aged dermal fibroblasts suppress the growth of primary melanoma while promoting metastatic dissemination to the lung (16). While these studies highlight the interplay between normal aged fibroblasts and cancer, how age impacts the phenotype and function of CAFs remains largely unexplored. In particular, it is unclear whether aged CAFs contribute to the immune landscape or metastasis in PDAC. Moreover, in tumor models where fibroblast function has been studied, it remains an open question whether reinvigorating aged stroma with more youthful fibroblastic properties can suppress or reverse the pro-tumorigenic effects observed *in vivo*.

In this study, using a syngeneic orthotopic PDAC mouse model, we demonstrate that organismal aging accelerates PDAC progression and enhances metastatic burden, findings that are not attributable to intrinsic tumor cell properties but instead implicate age-associated alterations in the tumor microenvironment. Through transcriptomic profiling and histological analysis, we identify that aging promotes a pro-metastatic stromal terrain driven by a fibroblast-driven program of ECM remodeling that contributes to epithelial-mesenchymal transition (EMT) and immune exclusion. Notably, we define a CAF-specific ECM gene signature that is significantly upregulated in aged human PDAC tumors, particularly within myofibroblastic CAFs, and correlates with adverse outcomes in older patients. To functionally test the impact of stromal aging, we employ heterochronic co-transplantation experiments, demonstrating that revitalization of the aged tumor microenvironment with young CAFs attenuates metastatic spread and restores immune infiltration, whereas introduction of aged CAFs into young hosts promotes immune suppression without significantly altering primary tumor growth. Overall, our findings establish aging CAFs as key architects of a pro-tumorigenic microenvironment and highlight their central role in mediating age-related PDAC aggressiveness. Moreover, they raise the provocative possibility that targeting age-associated changes in CAFs—either by fibroblastic rejuvenation or modulating the ECM—could mitigate the pro-tumorigenic effects of the aged stroma and enhance the efficacy of anti-cancer therapies. Our work establishes a new conceptual framework for understanding how aging shapes the tumor microenvironment in PDAC, opening a fertile avenue of investigation into age-specific stromal regulation, and raising compelling questions about the molecular mechanisms that mediate these effects—questions that our study now makes possible to ask.

## Results

### Age accelerates PDAC progression

To investigate whether organismal age impacts PDAC tumor progression or metastasis, we employed a syngeneic orthotopic mouse model. Murine KPC cells of the genotype *Pdx1-Cre; LSL-KRas^G12D/+^; LSL-Tp53^R172H/+^* derived from an autochthonous genetically engineered mouse model of PDAC were surgically implanted directly into the pancreata of C57BL/6 mice that were either young (∼2-months) or old (∼19-months). Four weeks following implantation, animals were sacrificed, and primary tumors were analyzed. We found that the primary tumors in the pancreas were significantly larger in the old mice (Fig. 1A). In KPC-derived tumors, cancer cells exhibit regional differences in proliferative capacity, where cells located in the densely vascularized peripheral regions display increased levels of Ki-67, a proliferation marker (17). Consistent with the increased tumor growth that we observed, primary PDAC tumors from the old animals had significantly higher numbers of proliferative cells in these peripheral regions relative to the tumors grown in young mice (Fig. 1B, C). We also observed a decrease in the extent of apoptosis in the old tumors, as measured by staining for Cleaved Caspase-3, an apoptosis marker (Fig. 1D,E). Young and old animals were also examined for macrometastases to the diaphragm, liver, and intestine. We observed an overall enhancement in the occurrence of macrometastases to any organ site, as well as to the specific organ sites (Fig. 1F, Supplementary Fig. S1A). Image-based quantification of metastatic burden in individual organs revealed a significant increase in the sizes of macrometastases in the old animals relative to the younger animals (Fig. 1G-I). Interestingly, the lung represented a notable exception to the age-associated metastatic phenotype, as no significant differences in metastatic tumor burden were observed between young and old animals. This assessment was based on p53 immunohistochemical staining, which specifically labels infiltrating KPC cells within the organ (Supplementary Fig. S1B). This is in contrast to a recent study that found that the lungs of older mice provide a more permissive niche for melanoma cancer cells to metastasize (18). Collectively, these findings indicate that PDAC progression is markedly accelerated in older organisms and suggest an age-dependent metastatic tropism influencing the dissemination of tumor cells to distant organs. Furthermore, given that the KPC cells utilized in both young and old mice were identical, the observed effects cannot be attributed to intrinsic tumor cell properties. Instead, these results implicate the age-associated alterations in the tumor microenvironment as the primary driving factor.

**Figure 1.**
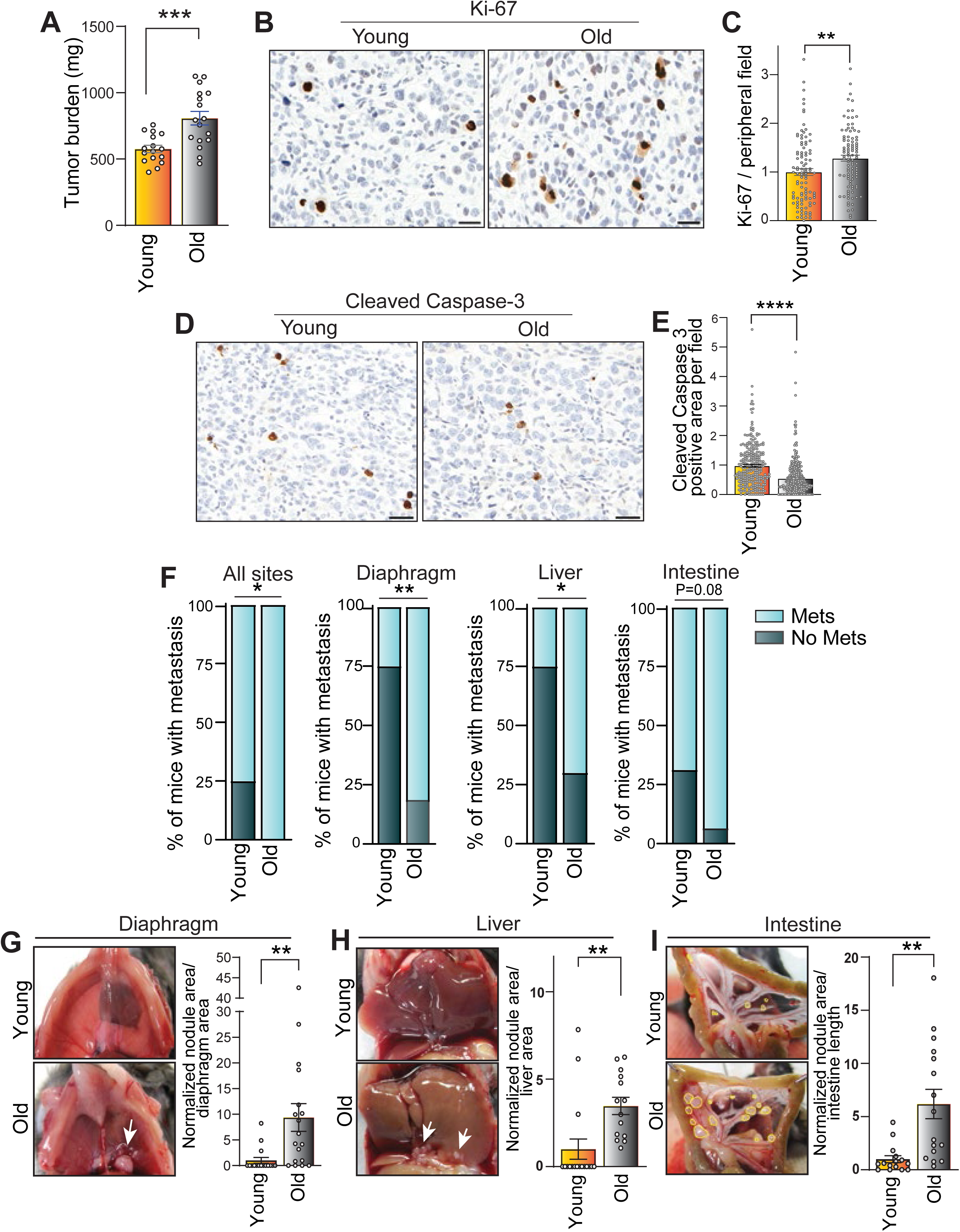
Aging accelerates tumor progression in a syngeneic orthotopic PDAC mouse model. **A,** Quantification of tumor weights of orthotopic PDAC tumors derived from young (n=16) and old (n=17) mice. Data are presented as mean ± SEM, unpaired Student *t* test, *** p ≤ 0.001. **B−C,** Immunohistochemical quantification of the proliferation marker Ki-67 in the periphery of orthotopic tumors, n=10 mice per group, ∼100 image fields from each group were analyzed; Scale bars, 100 μm, 20X magnification. **D−E,** Immunodetection of Cleaved Caspase-3 in orthotopic tumors grown in young and old mice; n = 10 mice per group. Scale bar 100 μm. Data are shown as mean ± SEM, unpaired Student *t* test, ** p ≤ 0.01, *** p ≤ 0.001, ****p < 0.0001**. F,** Graphs showing proportions of mice with KPC-derived primary tumors that exhibit macrometastases or no macrometastases to any organ (all sites, p = 0.04), diaphragm (p = 0.0016), liver (p = 0.014), and intestines (p = 0.08); n = 16 for young, n = 17 for old. p values were determined using two-tailed Fisher’s exact test. **G−I,** Multi-organ metastatic burden across various organ sites as indicated, accompanied by image-based quantitation. Bar plots display fold changes in nodular area between young and old tumor-bearing mice. The total area of tumor nodules was normalized to the organ total area (diaphragm and liver) or length (intestine). Data are presented as mean ± SEM, n = 16 for young mice, and n = 17 for old mice; 16 images from young mice and 17 images from old mice were analyzed for each organ. Outliers identified using Grubb’s test (α = 0.01) were excluded, unpaired Student *t* test; * p ≤ 0.05, ** p ≤ 0.01.

### Aging provokes a tumor-promoting stromal landscape in PDAC

To achieve a comprehensive characterization of age-associated differences in PDAC tumors, we conducted transcriptomic profiling through bulk RNA sequencing of tumors harvested from both young and old mice. Differential expression comparison of old versus young mice revealed a total of 546 differentially expressed genes (DEGs), with 300 genes upregulated and 246 genes downregulated (−0.5 ≤ Log_2_FC ≥ 0.5, FDR < 0.1) in tumors from older mice relative to their younger counterparts (Fig. 2A). In PDAC, diminished T cell infiltration is associated with poor patient prognosis, whereas increased CD8⁺ T cell infiltration has been positively correlated with improved patient survival (19–21). Consistent with this, gene set enrichment analysis (GSEA) of old versus young revealed a negative enrichment score (NES = −1.76, p < 0.05) for a gene set associated with the regulation of T cell-mediated immunity, indicating a decline in T cell functionality in tumors from older mice (Fig. 2B). Notably, expression levels of *Cd4* and *Cd8a*, markers of distinct T cell subsets, were downregulated in tumors from older mice, along with *Cd7*, a pan-T cell marker (Fig. 2A). Consistent with our transcriptomic findings, immunohistochemical evaluation confirmed a significant reduction in CD4^+^ and CD8^+^ T cells in tumors from older mice, suggesting an age-associated decline in T cell infiltration within the tumor microenvironment (Fig. 2C-D). In cancer, increased collagen deposition within tumors is often associated with reduced T cell infiltration, contributing to an immunosuppressive microenvironment. Dense collagen networks create physical barriers that impede the trafficking and infiltration of T cells into the tumor core. Moreover, ECM remodeling, driven by CAFs, can lead to the formation of a stiff, fibrotic stroma that further restricts immune cell access, acting as a structural impediment (22–26). To assess whether collagen deposition was more pronounced in the aged tumor microenvironment, we performed Masson’s trichrome staining on tumor sections, which revealed a significant accumulation of collagen in tumors from older mice compared to those from younger mice (Fig. 2E). These findings demonstrate that the aged PDAC microenvironment is characterized by reduced T cell infiltration and increased collagen deposition, which may contribute to the observed immune exclusion.

**Figure 2.**
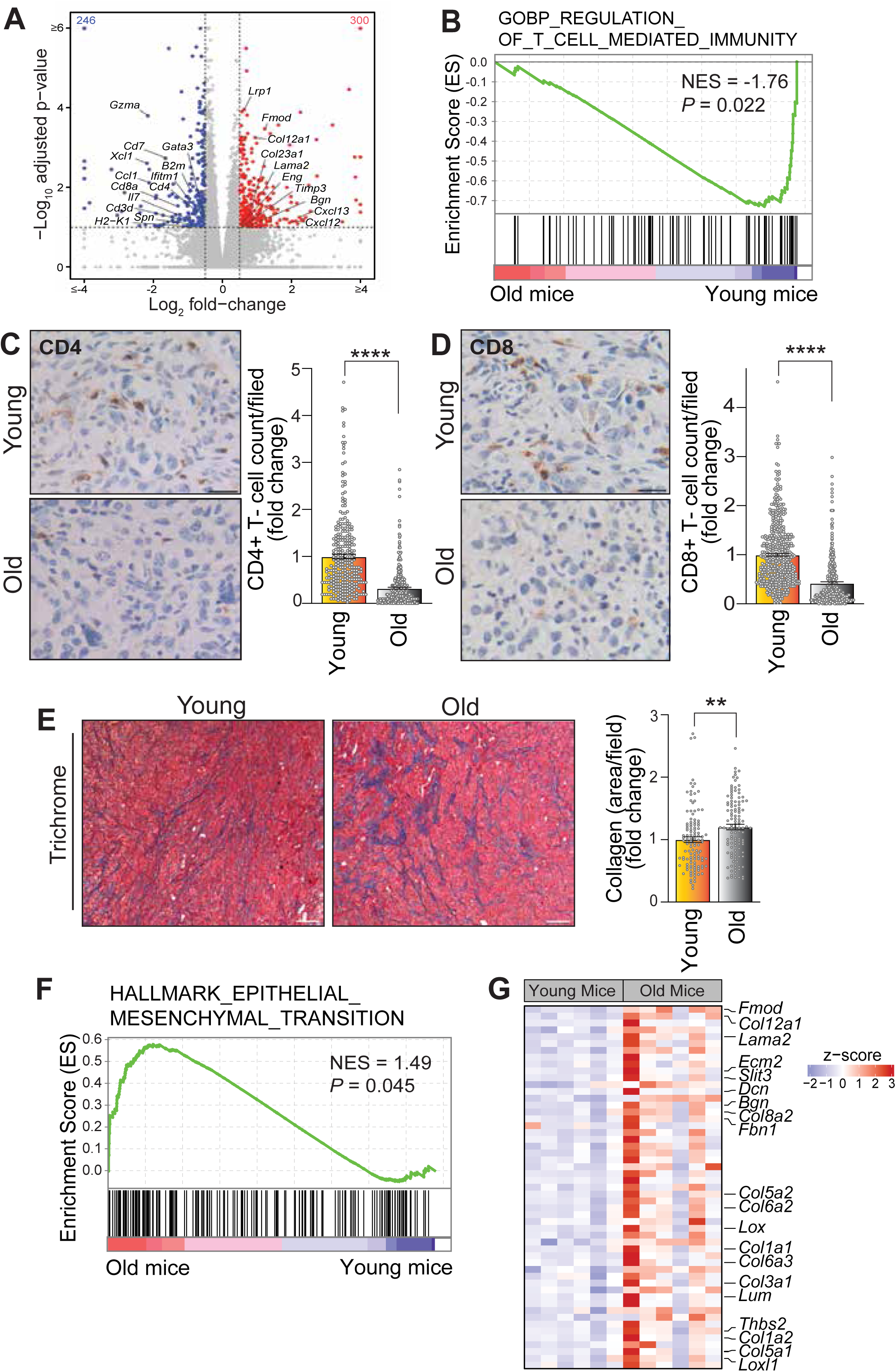
Age impacts the stromal landscape in PDAC tumors. **A,** Volcano plot showing differential gene expression comparison of orthotopic tumors extracted from old (n = 6) versus young mice (n = 6), with |FC| > 1.5 and FDR < 0.1. Blue dots indicate significantly downregulated genes (246 genes), and red dots indicate genes that were significantly upregulated (300 genes) in tumors grown in old mice. Representative significant genes related to T cell function and EMT are highlighted. **B,** Gene set enrichment analysis (GSEA) of bulk RNA-seq highlighting down-regulation of T-cell mediated immunity gene set in tumors derived from old versus young mice. NES, normalized enrichment score. **C−D,** Immunohistochemical staining for CD4⁺ and CD8⁺ T cells in tumor sections from young and old mice. For quantification, ∼300 image fields from young tumors (n = 10) and ∼350 image fields from old tumors (n = 10) were analyzed, Scale bar: 100 μm, 40X magnification. **E,** Masson’s trichrome staining of tumor sections was used to assess collagen deposition (blue). Bar plot displays fold difference in collagen-positive areas between tumors from young and old mice, ∼120 image fields (20X) from young tumors and ∼140 image fields (20X) from old tumors were analyzed. **F,** GSEA of epithelial-mesenchymal transition (EMT) gene set in tumors from young and old mice, NES, normalized enrichment score. **H,** Heat map of RNA-seq expression z-scores computed for ECM signature genes differentially expressed between the tumors from young and old mice.

In addition to the decline in T cell infiltration in tumors from older mice, we also found a significant positive enrichment of genes associated with epithelial-mesenchymal transition (EMT) (NES = 1.49, p < 0.05) in aged tumors (Fig. 2F). Given the well-established role of EMT in enhancing tumor cell invasiveness and metastatic potential, this finding provides further support for the accelerated metastatic progression observed in the older mice. We observed that of the 53 genes driving the tumor enrichment of the EMT signature in aged animals, 28 of them were predominantly involved in ECM organization, fibrosis, and/or tissue remodeling, processes that are integral to tumor progression, stromal interactions, and metastatic potential (Supplementary Fig. S2). These included genes encoding ECM structural proteins such as collagens, fibronectin-related proteins, proteoglycans, and laminins, as well as ECM-modulating factors, including lysyl oxidases and fibromodulin. Given their functional roles in shaping the tumor microenvironment, we curated this 28-gene signature as a molecular representation of the ECM alterations associated with aging in PDAC tumors (Fig. 2G). This signature serves as a valuable tool to characterize the fibrotic landscape of aged tumors and may provide insights into the mechanisms driving metastatic progression in older hosts. Moreover, our findings highlight a strong association between aging, ECM remodeling, and metastatic progression in PDAC, suggesting that age-related stromal alterations may play a critical role in driving tumor aggressiveness.

### Single-cell analysis reveals CAF-driven age-associated ECM remodeling in human PDAC

To determine whether the cellular processes enriched in tumors from older mice exhibit a similar age-dependent impact in human PDAC, we re-analyzed a single-cell RNA-sequencing (scRNA-seq) dataset from PDAC patients (27). UMAP clustering displayed distinct cell populations, which were annotated according to cell-specific marker genes described by Werba et al. (Fig. 3A and Supplementary Fig. S3A) (27). Importantly, our 28-gene ECM signature, identified from tumors in aged mice, showed a CAF-predominant expression pattern in human PDAC patients (Supplementary Fig. S3B). CAFs are known to be crucial to ECM remodeling, desmoplasia, and mechanical stiffness within the PDAC microenvironment, and these processes contribute to the aggressive and metastatic nature of PDAC. Therefore, to refine our gene signature and ensure specificity to CAFs, we excluded genes that exhibited non-specific expression across multiple cell types, as well as those with average Z-scores below 2 (e.g., *IRRC15, ABI3BP, LAMA1, LAMC1, COL4A1, FBLN5, NID2, COL4A2*, and *COMP*). Through this stringent filtering approach, we generated a 20-gene ECM signature with enhanced specificity to CAFs (Fig. 3B). Next, we examined whether this refined signature correlates with patient age. We observed a significant upregulation of the 20-gene ECM signature score in CAFs from older adult patients (>60 years) compared to younger adults (<60 years) (Fig. 3C). Further, UMAP-based re-clustering of the CAFs identified the two major CAF subtypes found in PDAC: myofibroblastic CAFs (myCAFs) and inflammatory CAFs (iCAFs) (Fig. 3D and Supplementary Fig. S4A). We found that myCAFs exhibited the highest ECM signature score among all analyzed cell populations (Supplementary Fig. S4B). Notably, the ECM signature score was significantly elevated in myCAFs from older patients, whereas in iCAFs, the signature score was reduced in older patients compared to younger ones (Fig. 3E). Previously, Werba et al. reported that while there were variations in CAF cell numbers across samples, the myCAF:iCAF ratio did not significantly differ between the Moffitt subtypes (basal and classical) (27). In our analysis, the overall proportion of myCAFs and iCAFs remained unchanged between younger (<60 years) and older (>60 years) individuals, indicating that the age-associated increase in the ECM signature is a cell type-specific phenomenon rather than a consequence of differences in CAF abundance across patients (Supplementary Fig. S4C). Next, we evaluated the prognostic significance of the individual genes comprising the 20-gene ECM signature by performing a hazard ratio analysis on the TCGA-PAAD dataset, stratified by age. In patients over 60 years, multiple genes (15 out of 20) — including *FMOD, COL12A1, LAMA2, BGN, FBN1, COL5A2, COL6A2, LOX, COL1A1, COL6A3, COL3A1, LUM, THBS2, COL1A2*, and *COL5A1*—exhibited a significantly elevated hazard ratio (HR > 1, p < 0.05), indicating an increased risk of adverse outcomes. In contrast, only a minority (4 out of 20) of these genes showed a significant association with poor prognosis in patients aged 60 or younger (Fig. 3F). Altogether, these findings indicate that age-related ECM remodeling in PDAC is primarily driven by myCAFs and correlates with poor prognosis in older patients, underscoring the potential of the 20-gene ECM signature as a prognostic biomarker and a tool for understanding stromal contributions to tumor progression.

**Figure 3.**
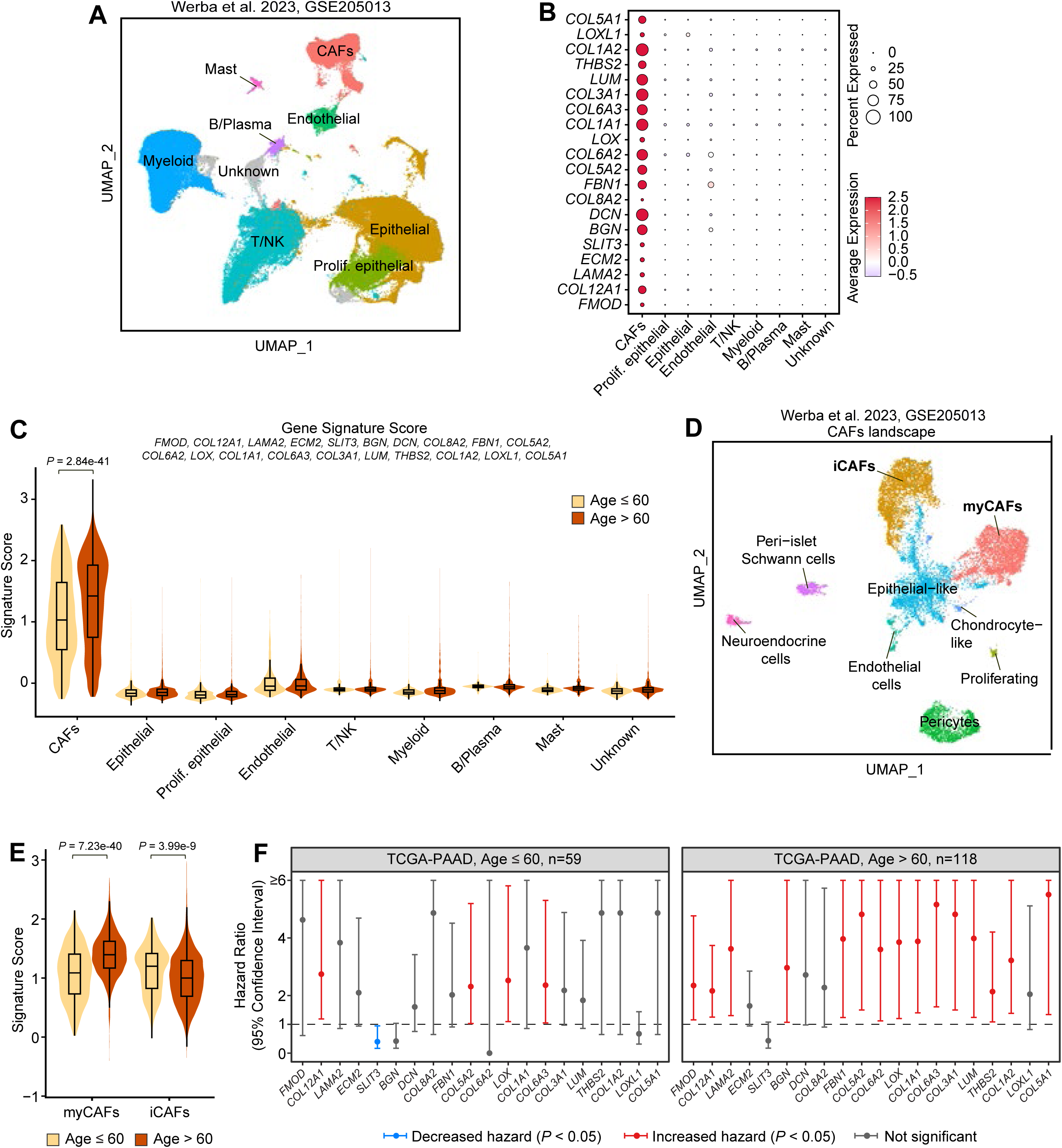
Aging promotes ECM remodeling in human PDAC. **A,** UMAP plot of major cell types obtained from re-analysis of human PDAC samples from Werba et al. (GSE205013). Cell types were identified using expression patterns of marker genes (see Supp. Fig 3A). **B,** Dot plot showing scaled expression levels in human PDAC cell types of ECM signature genes (originating from the EMT GSEA list) identified from bulk RNA-seq comparison of old versus young mice. Size of circles represent percent of cells expressing the genes and colors represent scaled expression levels. **C,** ECM signature score (based on average expression of the 20 genes), stratified by patient age (age ≤ 60 and age > 60) shows increased expression of the ECM signature in older patients (Wilcoxon Rank Sum test). **D,** UMAP of reclustering of the CAF cells in Werba et al. human PDAC dataset resolves CAF subtypes with distinct myCAF and iCAF clusters identified based on expression of marker genes. **E,** ECM 20-gene signature score in human PDAC myCAF and iCAF subtypes indicates that the ECM signature difference is more pronounced in myCAFs compared to iCAFs (Wilcoxon Rank Sum test). **F,** TCGA-PAAD patient cohort was divided into young (age ≤ 60, n=59) and old (age > 60, n=118), and hazard ratios were computed for high versus low expressors of our 20 ECM signature genes. Hazard ratio < 1 indicates decreased hazard (blue, *P* value <0.05), whereas hazard ratio >1 indicates lower survival (red, *P* value <0.05). Error bars represent 95% confidence intervals. Hazard ratios in grey are not significant.

### Reciprocal heterochronic transplantation reveals dominant anti-metastatic effects of the young microenvironment and immune-modulating roles of aged CAFs

Our findings underscore the role of the stroma in mediating age-dependent differences in the progression and metastatic potential of PDAC tumors. Given this, we next investigated whether invigorating the aged tumor microenvironment with young CAFs might reverse some of the observed age-related phenotypes. The heterochronic parabiosis model has been utilized as a tool to decipher the functional impact of systemic factors from young or old organisms on disease progression (28). Specifically, exposure of old mice to a young systemic environment elicits rejuvenation effects and immune-modulating functions (29, 30), while administration of systemic factors from old animals to young mice induces adverse effects such as impairments in neurogenesis and cognition (31). These and other findings highlight the critical impact that an aging microenvironment can have on tissue function and disease progression (28). To test these concepts in our model system, we employed a heterochronic co-transplantation approach where murine CAFs isolated from old or young tumor-bearing animals are orthotopically co-implanted with KPC cells into old or young mice. We first isolated murine CAFs from orthotopic PDAC tumors grown in young or old mice that were positive for both fibroblast-specific protein 1 (FSP1) and podoplanin (PDPN), established CAF markers (Supplementary Fig. S5A). We also verified that the isolated cells were negative for the epithelial marker E-Cadherin and positive for α-SMA, a well-established fibroblastic marker that all CAFs express *in vitro* (Supplementary Fig. S5B). In our first heterochronic co-implantation experiment, we co-injected either young or old CAFs together with KPC cells into the pancreata of old recipient mice (Fig. 4A). Weight of primary tumors and metastatic burden were quantified 28 days post-implantation. Notably, although revitalization of the old stroma with young CAFs did not significantly impact the weight of the primary tumors or the incidence of metastases to specific organs, there was a striking decrease in the metastatic tumor burden in the diaphragm, liver, and intestine (Fig. 4B-F and Supplementary Fig. S5C). We also observed that young CAFs had the capacity to significantly attenuate the enhanced collagen deposition associated with tumors in older mice, while concurrently promoting increased infiltration of both CD8⁺ and CD4⁺ T cells into the tumor microenvironment of aged hosts (Fig. 4G-I). These findings demonstrate that revitalization of the aged tumor stroma with young CAFs can mitigate key aspects of age-associated PDAC progression, including reducing metastatic tumor burden and restoring immune infiltration, despite having no significant effect on primary tumor growth. This highlights the potential for targeting stromal aging as a therapeutic strategy to counteract the pro-metastatic and immunosuppressive features of the aged tumor microenvironment.

**Figure 4.**
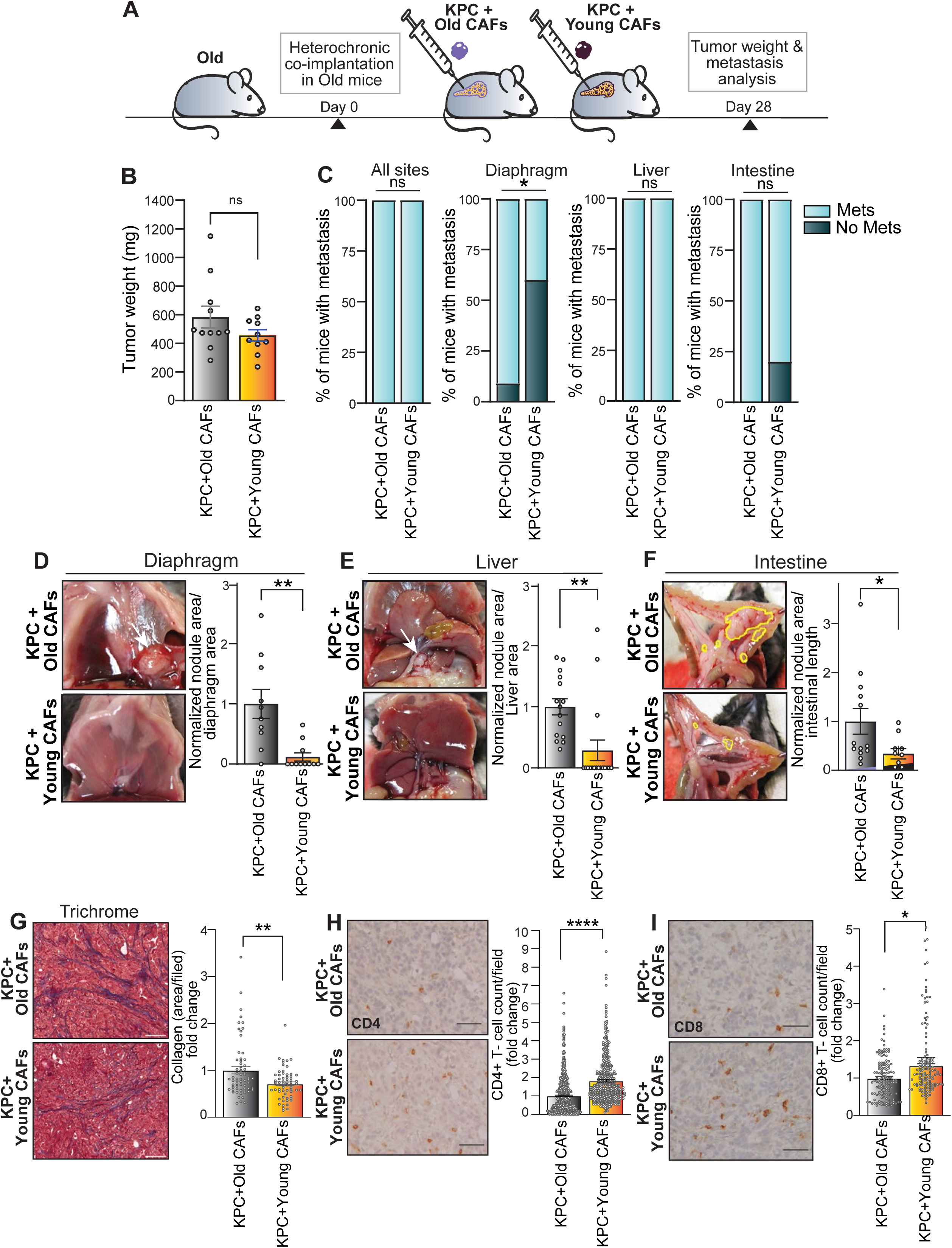
Revitalization of the old tumor stroma attenuates age-dependent acceleration of PDAC progression. **A,** Schematic of the heterochronic model in which KPC cells were orthotopically co-implanted with either young or old CAFs into the pancreata of aged mice. **B,** Graph displays tumor weights from orthotopic tumors derived in aged hosts following KPC cell co-implantation with old CAFs (KPC + Old CAFs, n = 11) or young CAFs (KPC + Young CAFs, n = 10). Data are presented as mean ± SEM, unpaired Student *t* test with Welch’s correction; n.s., non-significant. **C,** Proportions of aged mice that developed macrometastases or no macrometastases to any organ (all sites), diaphragm, liver, or intestines. KPC + Old CAFs (n = 11) or KPC + Young CAFs (n = 10); two-tailed Fisher’s exact test, p values are as indicated. **D−F** Multi-organ metastatic burden to various organ sites with quantitation. Bar plots indicate fold differences in metastatic nodule area between the two groups, i.e., KPC + Old CAFs (n = 11) versus KPC + Young CAFs (n = 10). The total area of tumor nodules was normalized to organ total area or length. Outliers identified using Grubb’s test (α = 0.01) were excluded. Data are presented as mean ± SEM, analyzed using unpaired Student *t* test, * p ≤ 0.05, ** p ≤ 0.01. **G,** Masson’s trichrome-stained tumor sections to visualize collagen deposition, and fold differences in collagen-positive areas were quantified, comparing KPC + Old CAFs (n = 11) versus KPC + Young CAFs (n = 10), ∼200 fields per group were analyzed at 20X magnification. **H−I**, Immunohistochemical staining for CD4⁺ and CD8⁺ T cells in tumor sections is shown. Bar plots indicate fold changes in CD4⁺ or CD8⁺ T cell counts comparing KPC + Old CAFs (n = 11) versus KPC + Young CAFs (n = 10), ∼400 image fields (40X) per group were analyzed. Scale bar: 100 μm. Data are represented as mean ± SEM; unpaired non-parametric Student *t* test*, \** p ≤ 0.05, ** p ≤ 0.01, **** p ≤ 0.0001, n.s., non-significant.

We next conducted a complementary heterochronic co-implantation experiment where young or old CAFs were orthotopically co-injected with KPC cells into young recipient mice (Fig. 5A). We found that reconstitution of the stroma with old CAFs in young mice did not lead to a significant increase in primary tumor growth or the incidence of metastases to distant organ sites (Fig. 5B,C and Supplementary Fig. S5D). Unexpectedly, metastatic burden across the analyzed organs remained unchanged, with the exception of a non-significant 2-fold increase in liver metastases (p = 0.0800) (Fig. 5D-F). This potential increase in metastatic propensity in the liver, driven by old CAFs in young mice, may represent another example of age-associated metastatic tropism in PDAC. We also did not observe changes in collagen deposition (Fig. 5G). Notably, augmentation of the PDAC tumor stroma with old CAFs in young mice led to a significant reduction in the numbers of intratumoral CD4⁺ and CD8⁺ T cells, suggesting an immunosuppressive effect driven by aged fibroblasts (Fig. 5H-I). These findings indicate that the endogenous tumor microenvironment in the young recipient mice may exert a dominant regulatory effect that counteracts the metastatic influence of old CAFs. Additionally, our data suggest that the old CAFs have the capacity to attenuate T-cell infiltration in the young stromal milieu, highlighting a potential immunosuppressive role of aged CAFs, independent of ECM remodeling, reinforcing the impact of stromal aging on the tumor immune microenvironment.

**Figure 5.**
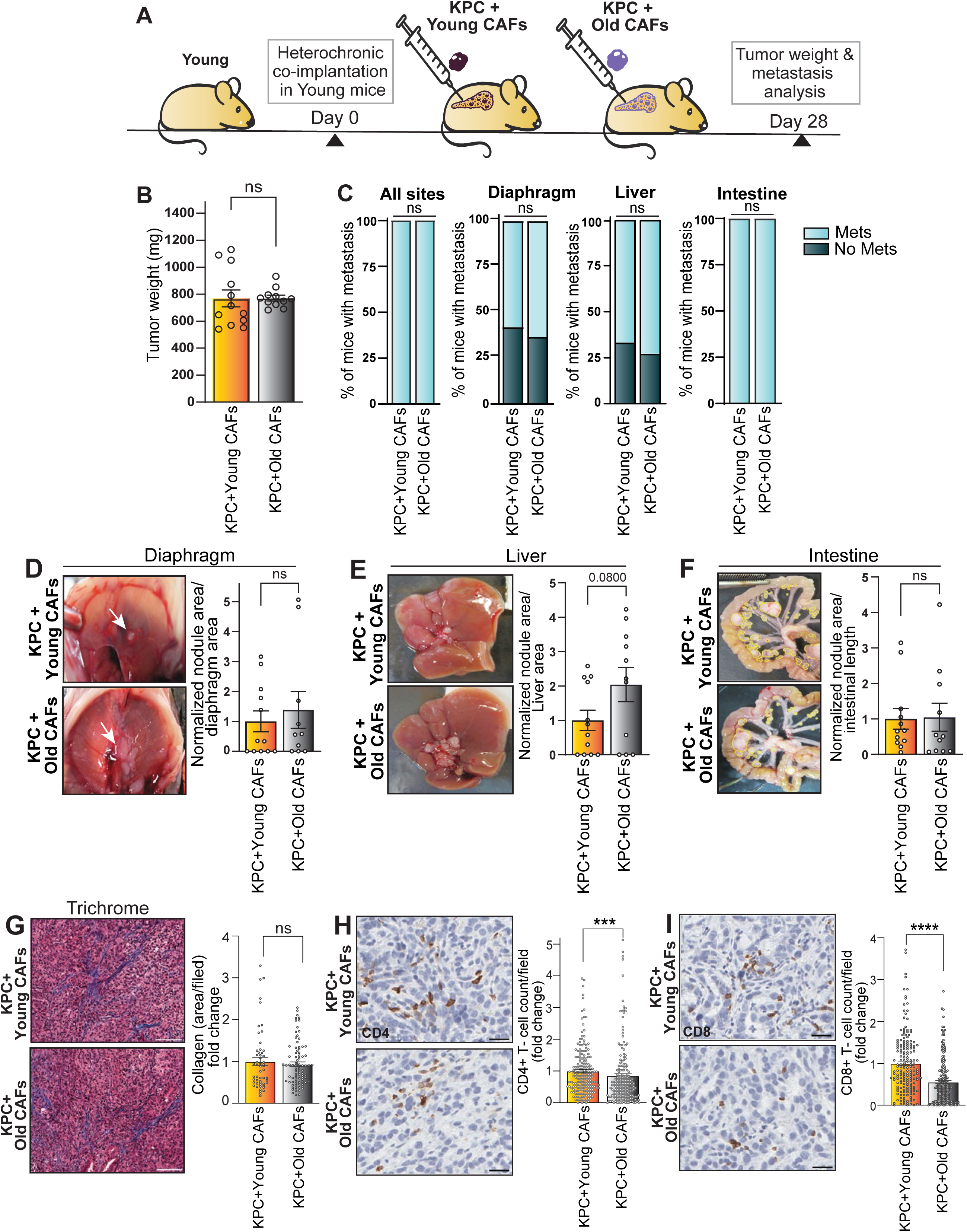
Tumor stroma of young hosts overrides aged CAF-driven pro-metastatic influence without abrogating inflammatory cues. **A,** Schematic of heterochronic model in which KPC cells were orthotopically co-implanted with either young or old CAFs into the pancreata of young mice. **B,** Graph displays tumor weights from orthotopic tumors derived in young hosts following KPC cell co-implantation with young CAFs (KPC + Young CAFs, n = 12) or old CAFs (KPC + Old CAFs, n = 11). Data are presented as mean ± SEM, unpaired Student *t* test with Welch’s correction; n.s., non-significant. **C,** Proportions of young mice that developed macrometastases or no macrometastases to any organ (all sites), diaphragm, liver, or intestines, statistical significance evaluated by using two-tailed Fisher’s exact test, n.s., non-significant. **D– F**, Representative images and quantification of metastatic burden to multiple organ sites. Bar plots indicate fold differences in metastatic nodule area between the two groups. Metastatic areas were normalized to total organ area or length. Data are presented as mean ± SEM, n = 12 for KPC + Young CAFs, n = 11 for KPC + Old CAFs, with outliers removed using Grubb’s test (α = 0.01). Statistical analysis performed using Student *t* test; n.s., non-significant. **G,** Masson’s trichrome-stained tumor sections are shown. Fold differences in collagen-positive areas between the groups were quantified, ∼200 fields per group at 20X magnification. n.s., non-significant. **H–I**, Immunohistochemical staining of CD4⁺ and CD8⁺ T cells in tumor sections from tumors derived in young mice, comparing KPC + Young CAFs versus KPC + Old CAFs. ∼400 fields (40X) per group were analyzed. Data are represented as mean ± SEM, p values were calculated by unpaired non-parametric Student *t* test; *p ≤ 0.05, **p ≤ 0.01.

### Discussion

Preclinical models of PDAC often overlook age-related physiological and pathological alterations, stalling the translational benefits to older adults who endure most of the disease burden. The aging tumor microenvironment can dramatically influence how cancer grows and spreads by modifying the extracellular matrix and immune landscape. In our study, we establish an age-relevant preclinical model where tumors grown orthotopically in the pancreata of old mice exhibit significantly enhanced proliferative activity, reduced apoptosis, and increased metastatic burden compared to tumors in younger mice. Our prior utilization of this particular model has demonstrated that orthotopic implantation of these KPC cells results in the development of sizable tumors that can metastasize to various organ sites in young mice (32–34); hence, the accelerated tumor burden we observed in aged hosts perhaps suggests earlier tumor initiation and dissemination driven by the age-modified tumor microenvironment. Our study advances the field of aging and pancreatic cancer by providing direct *in vivo* evidence that stromal aging acts as an accelerator of tumor progression. Moreover, these findings highlight the importance of age-stratified preclinical models to pave the way for precision interventions by targeting the unique biology of elderly PDAC patients.

Our transcriptomic analysis provides mechanistic insights into age-related discrepancies driving tumor progression, primarily through processes involving immune evasion, epithelial-mesenchymal transition, and stromal remodeling. The reduction in the numbers of intratumoral CD4^+^ and CD8^+^ T cells is likely indicative of restricted T-cell infiltration into tumors from old mice, consistent with the immunologically ‘cold’ nature of PDAC as reported using single-cell approaches (35). In previous studies, aging-induced alterations in the lungs were mediated via IL-17-γδT-neutrophil signaling axis, which suppresses CD8^+^ T cell function and promotes tumor metastasis (36). In PDAC, the reduction of CD4^+^ and CD8^+^ T cells likely stems from stromal aging, where excessive collagen deposition around tumors impedes T-cell infiltration. Furthermore, we identified a gene signature indicative of T cell–mediated immune dysfunction within aged tumors, with multiple components converging on pathways known to drive PDAC pathogenesis and immune evasion in humans. For instance, the coordinated downregulation of MHC-I components (B2M, TAP2, H2/MHC-I variants) in tumors from aged mice suggests impairments in antigen presentation resulting from the destabilization of MHC-I complex and reduced tumor antigen display. Such defects exacerbate the intrinsically low mutational burden in PDAC, leading to even more restricted antigen presentation (37). Likewise, tumor intrinsic factor Progranulin (PGRN) attenuates T-cell infiltration by destabilizing the MHC-I complex, an effect linked to poor overall survival in PDAC (38). These findings from our study and others suggest that factors secreted by the aged tumor stroma may disrupt the MHC-I complex, potentially contributing to the escape from CD8⁺ T-cell recognition. Next, we observed a paradoxical decrease in FOXP3 expression in aged tumors. Although overexpression of FOXP3 in PDAC typically correlates with poor prognosis through recruitment of immunosuppressive Tregs, its downregulation in older tumors may instead signal Treg exhaustion rather than complete loss of immunosuppression. This is supported by the continued presence of residual FOXP3^+^ cells within the stroma, in addition to dense stromal collagen, both of which sustain an immunosuppressive environment (39, 40). Overall, these findings entail impaired antigen presentation, altered Treg signaling, and collagen driven barriers that act together to promote immune evasion and metastatic spread. These observations emphasize the need for combined therapeutic strategies targeting both antigen presentation restoration (via FAK inhibition) and stromal collagen (via LOX inhibition) to counteract age-driven immunosuppression in pancreatic cancer (41, 42).

We identified a conserved age-related ECM-gene signature across mouse and human PDAC. Several genes from the ECM signature appear to link stromal aging to immunosuppression. Notably, the ECM-gene signature score was significantly elevated in myCAFs from older patients compared to younger ones while being reduced in iCAFs from older patients. Numerous genes from this signature, including LOXL1, LOX, THBS2, COL1A1/COL1A2 exhibit functional roles in PDAC pathogenesis. For example, lysyl oxidases (LOX/LOXL1) are known to enhance ECM stiffness and create an immunosuppressive niche in human PDAC by modulating tumor-infiltrating lymphocytes (TILs) and immune checkpoints (43). LOX inhibitors have been shown to alleviate fibrotic ECM and overcome chemoresistance in triple-negative breast cancer (44). Given the plausible role of LOXs in tumor immune evasion, combining LOX inhibitors with immune checkpoint blockade may help to overcome ECM barrier effects and restore T-cell trafficking within aged tumors. Another gene from the ECM signature−THBS2 combined with CA 19-9 serves as a diagnostic biomarker for early detection of human PDAC (45). THBS2 remodels the PDAC microenvironment via CD47-mediated signaling and promotes immune evasion. Knockdown of THBS2 reduces CD47 expression, leading to decreased proliferation, migration, and invasion of PDAC cells (46). Similarly, COL1A1/2 genes encode distinct oncogenic homotrimers that regulate PDAC microbiome and promote immune evasion through MDSC recruitment and T-cell exclusion. Disrupting cancer-specific COL1 reduces desmoplasia, depletes MDSCs, enhances T-cell infiltration, and increases sensitivity to anti-PD-1 immunotherapy (47). Although aging-specific roles for these genes remain unexplored, our data suggest that coordinated overexpression of ECM gene signature in aged CAFs creates a pro-tumorigenic niche, underlining the need for age-stratified therapeutic testing in preclinical models.

myCAFs are the primary producers of extracellular matrix in pancreatic cancer, secreting collagens, fibronectin, and matricellular proteins that stiffen the tumor stroma. The matrix supports tissue architecture, cell migration, and intercellular communication. In our study, the prognostic power of ECM gene signature derived from aged mice aligns with clinical data and predicts an increased risk of adverse outcomes in older PDAC patients. To elucidate how age-related stromal alterations contribute to tumor progression, we utilized heterochronic transplantation, allowing us to dissect whether invigorating the old tumor stroma could reverse age-dependent tumor phenotypes. Interestingly, the resuscitation of aged stroma with young CAFs counteracted aged-related tumor characteristics, including multi-organ metastasis, collagen deposition, and CD4⁺/CD8⁺ T cell infiltration into the tumor microenvironment. These findings are consistent with previous heterochronic studies where young systemic factors rejuvenated the aged tissues by altering transcriptional profiles and cellular senescence associated with aging (48, 49). However, the interaction between old CAFs and the young host stroma did not fully recapitulate the pro-tumorigenic environment observed in aged hosts, suggesting that young mice exert robust compensatory mechanisms, such as enhanced immune surveillance and/or more efficient DNA repair, which offset the pro-tumorigenic effects of old CAFs.

The primary unanswered question in the field is which underlying mechanisms can provide effective personalized therapies for elderly PDAC patients. Overall, our findings highlight the potential of targeting stromal aging for therapeutic benefits. For instance, re-engineering the tumor microenvironment through stromal normalization or a combination of antifibrotic drugs such as PXS-5505 with immunotherapy may reduce fibrosis and improve survival in elderly PDAC (42). The anti-fibrotic agent, Halofuginone is known to suppress collagen synthesis by targeting TGF-β signaling, also promotes immune infiltration and drug delivery through stromal normalization (50, 51). Similarly, targeting focal adhesion kinase (FAK) with anti-fibrotic agent VS-4718 limits PDAC progression and renders tumors more responsive to immunotherapy (52). Similarly, Nintedanib mediated inhibition of fibrogenic signaling (PDGFR, FGFR, VEGFR) may improve immune infiltration and enhance antitumor responses in the elderly population (53). However, challenges such as stromal complexity, cellular heterogeneity, and unique collagen dynamics associated with aging must be considered when targeting the tumor stroma. Future studies should investigate more personalized strategies for targeting fibrosis-dependent tumors guided by stromal profiling using tools such as our age-dependent ECM signature. Additionally, reprogramming the stroma using factors derived from young CAFs or heterochronic models to revitalize aged stroma could open new pathways for therapeutic advancement.

## Acknowledgments

We are grateful to members of the Commisso laboratory for their constructive comments. We thank Dr. Peter Adams and his laboratory for helpful discussions. We acknowledge Dr. Deepika Bhullar for contributions to the initial phase of this project, including performing early experiments that informed the development of this study. This work was supported in part by NIH grant R21AG087544 to C.C. We thank Dr. Robert Vonderheide for KPC4662 cells. We thank the Sanford Burnham Prebys Histology Core, Genomics Core, Bioinformatics Core, and Animal Facility. Sanford Burnham Prebys core services are supported by NCI Cancer Center Support grant P30CA030199.

## Author contributions

P.G. Data curation, formal analysis, investigation, visualization, methodology, writing–original draft, and project administration. L.L, Y.Z. K.D.P, S.M., and C.G. Investigation, data analysis, and data curation. R.M. Bioinformatics investigation, formal analysis, data curation, methodology, writing–review and editing. C.C. Conceptualization, data analysis, data interpretation, resources, supervision, funding acquisition, writing–original draft, writing–review, editing.

## Methods

### Cell culture and CAF isolation

KPC4662 cells were derived from a Kras^LSL-G12D/+^;Trp53^LSL-R172H/+^;Pdx1-Cre (KPC) female mouse and were kindly provided by Dr. Robert Vonderheide (University of Pennsylvania)(54). KPC cells were cultured in DMEM complete medium (high glucose, L-glutamine DMEM) containing 10% FBS, 100 Units/mL penicillin/streptomycin, and 20mM HEPES. Murine CAFs were isolated from PDAC tumors generated by orthotopic implantation of KPC4662 cells into the pancreata of either young (∼2-months) or old (18- to 21-months) female mice. Tumors were harvested, finely minced into ∼2 mm fragments using sterile razor blades, and rinsed with phosphate-buffered saline (PBS). Several tumor pieces were then plated in 10 cm culture dish containing 5 mL of complete DMEM supplemented with 100 µg/mL gentamycin and incubated at 37 °C under standard culture conditions. After 3 days, the tumor fragments were removed, and based on morphological evaluation, spindle-shaped fibroblast clusters located away from tumor cell colonies were digested in situ using 0.25% trypsin. Trypsinized cells were then transferred to 24-well plates for expansion. CAF lines were maintained in complete DMEM medium. Primary CAFs were transduced with a retroviral vector encoding the SV40 T antigen. Retrovirus was produced by co-transfecting HEK293T cells with the pBabe-zeo-SV40 vector and the pCL-Ampho packaging plasmid using Lipofectamine 2000 (Invitrogen, Cat# 11668019). Viral supernatants were collected 48 hours post-transfection, filtered through 0.45 µm syringe filters, and stored at −80 °C for future use. CAFs from young and old tumors were then transduced with the retrovirus. The fibroblast identity of the immortalized CAFs was confirmed by flow cytometric analysis and immunoblotting. For the flow cytometry identity validation, purified murine CAFs derived from young or old tumors were pre-incubated with blocking anti-CD16/32 antibody at 4 °C for 15 min and then stained with CoraLite®Plus (CL) 488-conjugated anti-FSP-1 (Proteintech, Cat#CL488-16105) and APC/Cy7-conjugated anti-PDPN (BioLegend, clone 8.1.1) antibodies at 4 °C for 30 min, followed by flow cytometric analysis. For the immunoblotting identity validation, cell lysates from KPC4662 and murine CAFs were prepared in RIPA buffer (10mM Tris-HCl [pH 8.0], 150mM NaCl, 1% sodium deoxycholate, 0.1% SDS, 1% Triton X-100) supplemented with 1x protease inhibitor cocktail and 1x PhosSTOP. Lysates were then centrifuged at 18,000g for 15 min at 4 °C, and protein concentrations were determined using a BCA Protein Assay Kit. Protein samples were separated by SDS-PAGE electrophoresis and then transferred to nitrocellulose membranes using Trans-Blot Turbo Transfer System (Bio-Rad), followed by incubation with primary antibodies: rabbit α-SMA (CST, 1:1000), rabbit E-cadherin (CST, 1:1000), and mouse β-Actin (Sigma, A2228, 1:5000). After staining with IRDye 680RD or 800CW secondary antibody (LI-COR), immunoblots were detected using Odyssey CLx imager (LI-COR).

### Animal experiments

C57BL/6 mice were purchased from Charles River Laboratories and acclimatized for a minimum of four days before experimentation. All animal procedures were performed at the Sanford Burnham Prebys Medical Discovery Institute animal facility, in compliance with protocols approved by the Institutional Animal Care and Use Committee (IACUC), ensuring adherence to ethical guidelines. Young (∼2-months, n = 16) and old (18- to 21-months, n = 16) female C57B/L6 mice were utilized for all orthotopic surgeries. Approximately 2.5 × 10^4^ KPC4662 cells were resuspended in PBS with 50% matrigel (Corning) and injected directly into the pancreata of C57BL/6 mice. Tumors were allowed to develop for 27 days post-surgery, after which mice were euthanized, tumors were harvested, and tumor weights were quantified. Tumor tissues were fixed in 10% neutral-buffered formalin for downstream immunohistochemical analysis. Macrometastases in the diaphragm, liver, and intestine were visually inspected, photographically documented, and recorded. For metastatic burden quantification, the nodule area normalized to the corresponding organ’s total area or length was determined using ImageJ software.

For the heterochronic co-transplantation model, a co-injection mixture of young or old murine CAFs combined with KPC4662 cells was prepared in a 1:3 ratio (2.5 × 10⁴ KPC4662 : 7.5 × 10⁴ CAFs). The cells were resuspended in PBS with 50% matrigel and surgically implanted into the pancreata of either young or old recipient mice. In the first heterochronic cohort, young CAFs + KPC4662 or old CAFs + KPC4662 were co-injected into the pancreata of old recipients (n = 10 and n = 11, respectively). In the second cohort, the same combinations were injected into young recipients (n = 12 and n = 11, respectively). All surgical procedures were performed under anesthesia, and animals were maintained on a heating pad until full recovery. Post-operative analgesia was provided via a single subcutaneous injection of extended-release buprenorphine (1.3 mg/mL) to minimize discomfort. Tumors were harvested 28 days post-implantation for weight measurement and histological processing. Macrometastatic lesions in the diaphragm, liver, and intestine were visually assessed, photo-documented, and quantified as described above using ImageJ.

### RNA-seq library preparation

Total RNA was isolated from tumor pieces (n=6 for young and old) using the PureLink RNA Mini Kit (Invitrogen, Cat# 12183025,) according to the manufacturer’s instructions. Isolated RNAs were quality-controlled using Agilent 4200 TapeStation system (Agilent Technologies) and only preparations with RIN > 8 were used to prepare cDNA libraries. RNA-seq libraries were built and sequenced as previously described (55). Briefly, PolyA RNA was isolated using the NEBNext Poly(A) mRNA Magnetic Isolation Module, and bar-coded libraries were constructed using the NEBNext Ultra Directional RNA Library Prep Kit for Illumina (NEB). Libraries were pooled and sequenced from a single end (175) using an Illumina NextSeq 500 system with the High Output V2 kit (Illumina). RNA-seq samples were sequenced at a depth of 18–25 million reads.

### RNA-seq data analysis

RNA-seq data processing and analysis was performed as previously described (55). Briefly, Illumina Truseq adapters and polyA/polyT sequences were trimmed using Cutadapt v2.3. Trimmed reads were aligned to mouse genome version mm10 and Ensembl gene annotations v84 using STAR version 2.7.0d_0221 (56) and ENCODE long RNA-seq pipeline parameters (https://github.com/ENCODE-DCC/long-rna-seq-pipeline). RSEM v1.3.1 (57) was used to obtain gene level estimated counts and transcripts per million. FastQC v0.11.5 (https://www.bioinformatics.babra ham.ac.uk/projects/fastqc/) and MultiQC v1.8 (58) were used to assess quality of trimmed raw reads and alignment to genome and transcriptome. Genes with RSEM estimated counts ≥5 times the total number of samples were retained for differential expression analysis. Differential expression comparison was performed using Wald test implemented in DESeq2 v1.22.2 (59). Genes with a Benjamini– Hochberg-corrected P value of < 0.1 and log_2_ fold-change of ≥0.5 or ≤-0.5 were defined as significant. Genes were ranked using a metric computed by multiplying log_2_ fold-changes and -log_10_ of adjusted p-values. Pre-ranked GSEA was performed using GSEA v4.3.3 (60) and MSigDB mouse gene set collections of hallmark and m5 GOBP version 2024.1 (61).

### Re-analysis of Werba et al. PDAC scRNA-seq

Processed human PDAC patient scRNA-seq sample data (count matrices) from Werba et al. (27) were downloaded from NCBI GEO under accession GSE205013. Data integration and analysis were performed using Seurat version 4.0.3 (62) in R version 4.0.2. Raw single-cell data were converted to Seurat object using *CreateSeuratObject(min.cells = 5, min.features = 500)*. Doublets were determined using scDblFinder (63)[PMID: 35814628]. Cells with <15% mitochondrial reads, <1% erythroid (HBA1, HBA2, and HBB) reads, and scDblFinder singlets were retained for downstream analysis. Seurat objects for the 27 patient samples were merged, data was normalized using *NormalizeData,* and top 2000 highly variable genes were determined using *FindVariableFeatures*. Data was scaled using *ScaleData* and PCA was run using *RunPCA* considering only the variable genes. Data for 27 patient samples were integrated using function *RunHarmony*. Cells were clustered using *RunUMAP*, *FindNeighbors*, and *FindClusters(resolution = 0.3)* with number of principal components (PCs) accounting for 90% of variance in the data. Cell clusters were annotated based on expression of marker genes described in Werba et al. We computed per cell score for our 20-gene ECM signature using *AddModuleScore* function. Comparison of this 20-gene ECM signature score was performed using *wilcox.test* function in R.

For CAF reclustering, normalized gene expression variances in CAF cells were computed using var function in R. Genes with variance >0.5 and mean expression <3.0 across all CAF cells were considered as highly variable and used for downstream analysis. CAF reclustering analysis was performed as described above for all cell types (*ScaleData*, *RunPCA*, *RunHarmony*, *RunUMAP*, *FindNeighbors*, *FindClusters*) with the exception of “resolution = 0.2” parameter in *FindClusters* step.

### TCGA-PAAD hazard ratios

Hazard ratios for each of our 20-gene ECM signature genes were computed as previously described (64). Briefly, gene expression and sample/patient information for TCGA PanCancer Atlas PAAD project were downloaded from cBioPortal (65). Hazard ratios were computed in R v4.4.1 using *survival* v3.7-0, *survminer* v0.5.0, and *maxstat* v0.7-25 packages. Optimal cutpoints for classifying patient samples as high or low expressors of genes of interest were determined using *surv_cutpoint* and *surv_categorize* functions. Hazard ratios, 95% confidence intervals, and p-values were computed using *coxph* function with overall survival (in months) and high versus low expressor patient sample classifications as described above.

### Immunohistochemistry

Formalin-fixed, paraffin-embedded (FFPE) tumor tissues were sectioned at 5 µm thickness. Sections were then deparaffinized, and antigen retrieval was performed by microwave heating in 0.01 M citrate buffer (pH 6.0). Endogenous peroxidase activity was quenched by incubating slides in 1.5% hydrogen peroxide for 30 minutes. For intracellular antigens, tissue sections were permeabilized with 0.1% Triton X-100 in TBS containing 0.1% Tween-20 (TBST) for 10 minutes. Following permeabilization, tumor sections were blocked with 10% goat serum plus 1% BSA for 1h at room temperature and incubated with primary antibodies diluted in 1% BSA in TBST overnight at 4 °C. After washing with TBST, slides were incubated with biotinylated secondary antibodies either goat anti-rabbit or goat anti-rat (Vector Laboratories; BA-1000, BA-9401) for 75 minutes at room temperature. The staining signal was detected using HRP-conjugated ABC detection system (Vector Laboratories, PK-6100) and DAB HRP Substrate Kit (Vector Laboratories, SK-4100). Sections were counterstained with hematoxylin, dehydrated, and mounted using Permount mounting medium (Fisher Scientific, SP15-100). The primary antibodies utilized were: rabbit anti-Ki-67 (Abcam, ab16667, 1:200), rabbit anti-CD8 (Abcam ab209775, 1:1500), rabbit anti-CD4 (Abcam ab183685, 1:500), rabbit anti-cleaved caspase 3 (CST 9664S, 1:600). Slides were imaged using Aperio AT2 digital scanner (Leica) or a bright-field Olympus CX-31 microscope equipped with INFINITY camera and capture software (Lumenera). Quantitative image analysis was performed using ImageScope software (Leica) or ImageJ (FIJI).

### Trichrome and H&E staining

Trichrome staining of the FFPE tumor sections was done using NovaUltra Masson Trichrome stain kit (Cat. No. IW-3006, IHC World). Briefly, tissue sections were deparaffinized and incubated in Bouin’s solution for 1hr at 56°C. After rinsing with water, slides were incubated in Weigert’s iron hematoxylin solution for 10 min. Afterward, tissue sections were stained with Biebrich scarlet acid fuchsin solution for 5 min. Tissue sections were incubated in Phosphotungstic/Phosphomolybdic (PP) Acid Solution for 15 min or until the collagen is no longer red followed by aniline blue staining for 20 min. After rinsing with distilled water, tissue sections were differentiated in acetic acid solution and after dehydrating, mounted with coverslips using Permount mounting medium. H&E staining of FFPE tumor samples were performed by Sanford Burnham Prebys Medical Discovery Institute Histology Core facility. Images were captured using a bright field microscope installed with INFINITY camera (Lumenera). For quantification, the collagen area per field of trichrome staining was determined using Fiji (ImageJ) software.

### Statistical analysis

All statistical analyses and graphs were made using GraphPad Prism software (version 10.4.1). Results are presented as mean ± standard error of the mean (SEM), unless otherwise stated. Comparisons between two groups were made using unpaired two-tailed Student *t* tests, with Welch’s correction applied when variances were unequal. For metastasis incidence, Fisher’s exact test (two-tailed) was used to assess statistical significance (recommended for small sample sizes to ensure exact p-values). Outliers in tumor weights and metastatic burden datasets were identified and excluded using Grubb’s test (α = 0.01). All experiments were independently performed at least three times. Data met the assumptions required for the statistical tests applied, with normality and variances formally assessed. p value less than 0.05 was considered statistically significant. Significance levels are indicated as: *p < 0.05,**p < 0.01, ***p < 0.001, ****p < 0.0001.

---

**Supplementary Figure 1.**
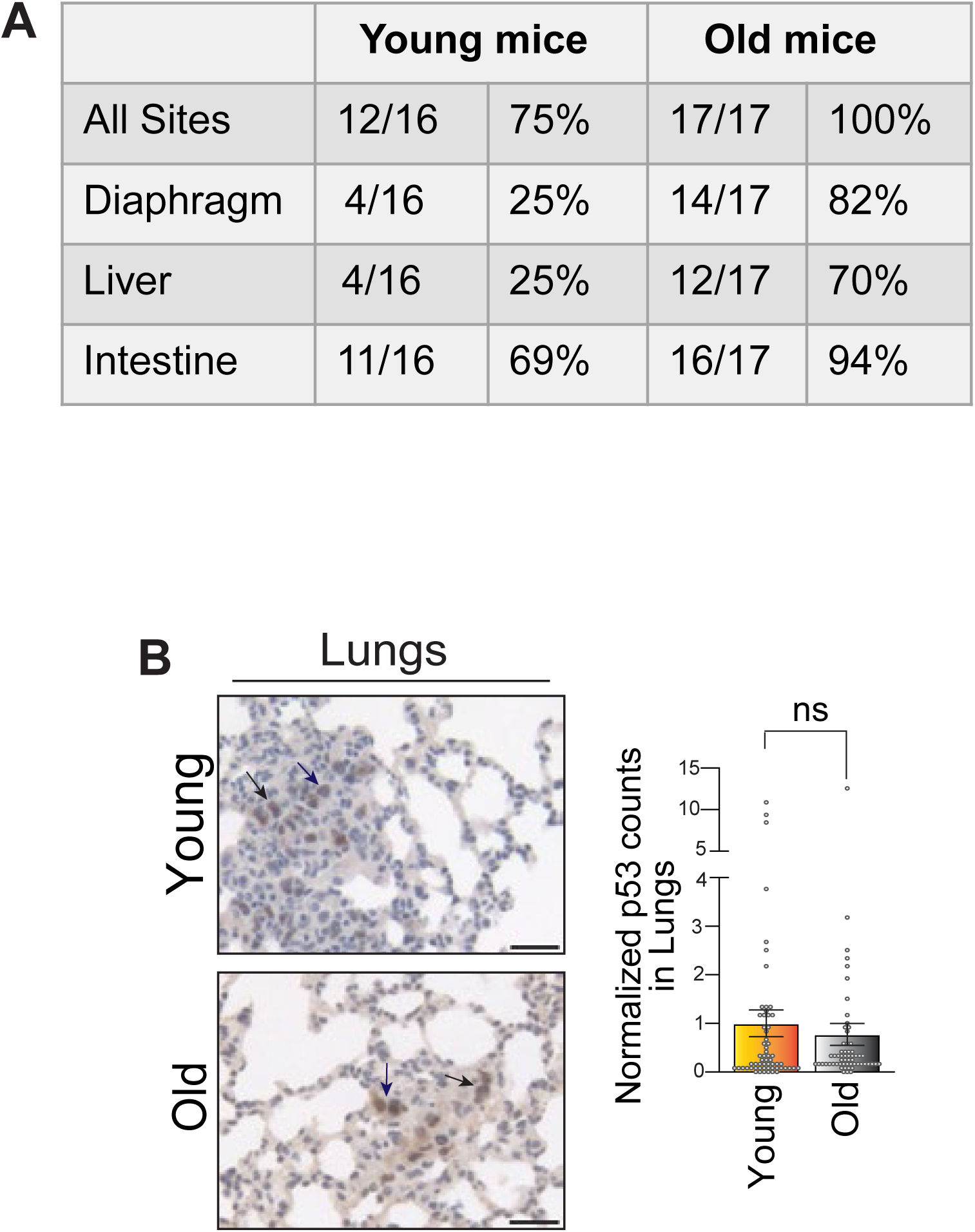
Metastatic incidence is influenced by host age. **A,** Table summarizing percentages of mice that presented with metastases across different organ sites; n = 16 or n = 17 mice per group. **B,** Immunodetection of p53 in lung tissues from young and old orthotopic tumor-bearing mice; graph shows fold change in p53-positive cell counts between young and old mice. n = 10 mice per group. Scale bar 100 μm. Data are represented as mean ± SEM, unpaired non-parametric Student *t* test, n.s., non-significant.

**Supplementary Figure 2.**
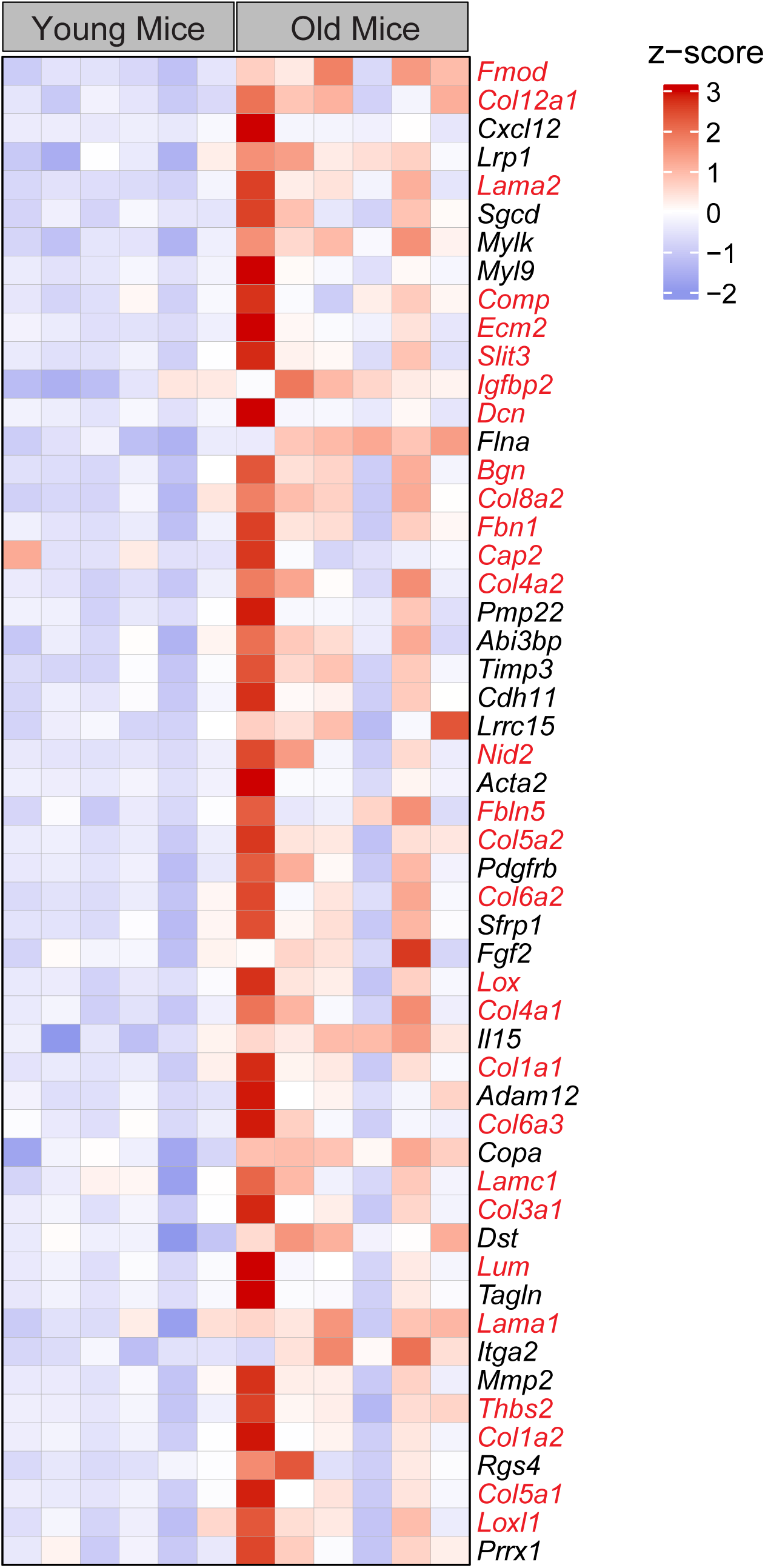
Age-associated enrichment of ECM gene signature in mouse PDAC. Heatmap showing scaled expression levels (z-scores) of core-enriched genes from pre-ranked GSEA analysis of old versus young mice. Genes highlighted in red were selected as our ECM signature gene set, which were further characterized in this study.

**Supplementary Figure 3.**
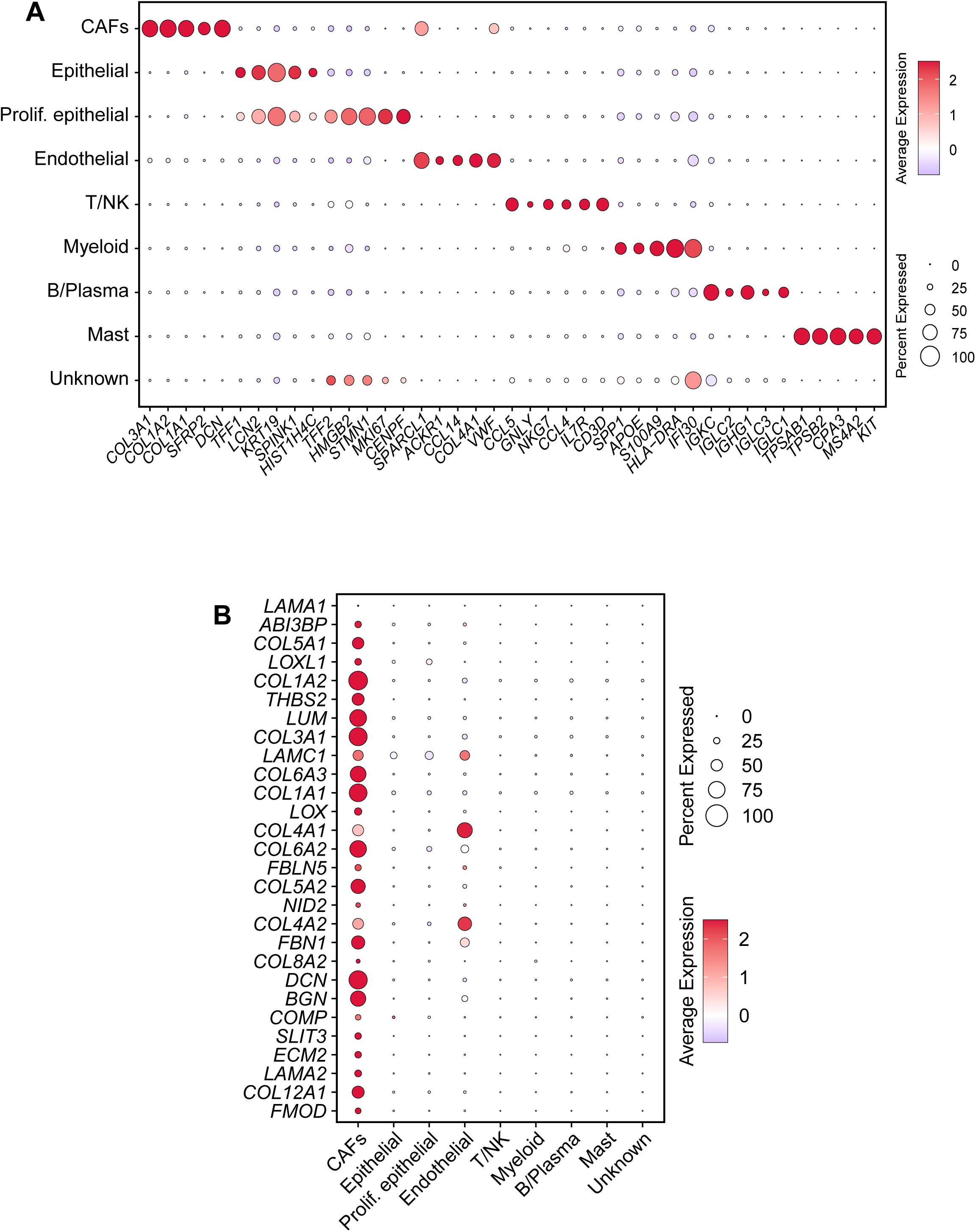
CAF-driven ECM gene signature expression in human PDAC. **A,** Dotplot showing scaled expression levels of marker genes to identify major cell types in re-clustering of human PDAC patient datasets from *Werba et al.* Marker genes were adopted from *Werba et al.* **B,** Dotplot showing scaled expression of our full 28 gene ECM signature identified from bulk RNA-seq comparison of old versus young mice.

**Supplementary Figure 4.**
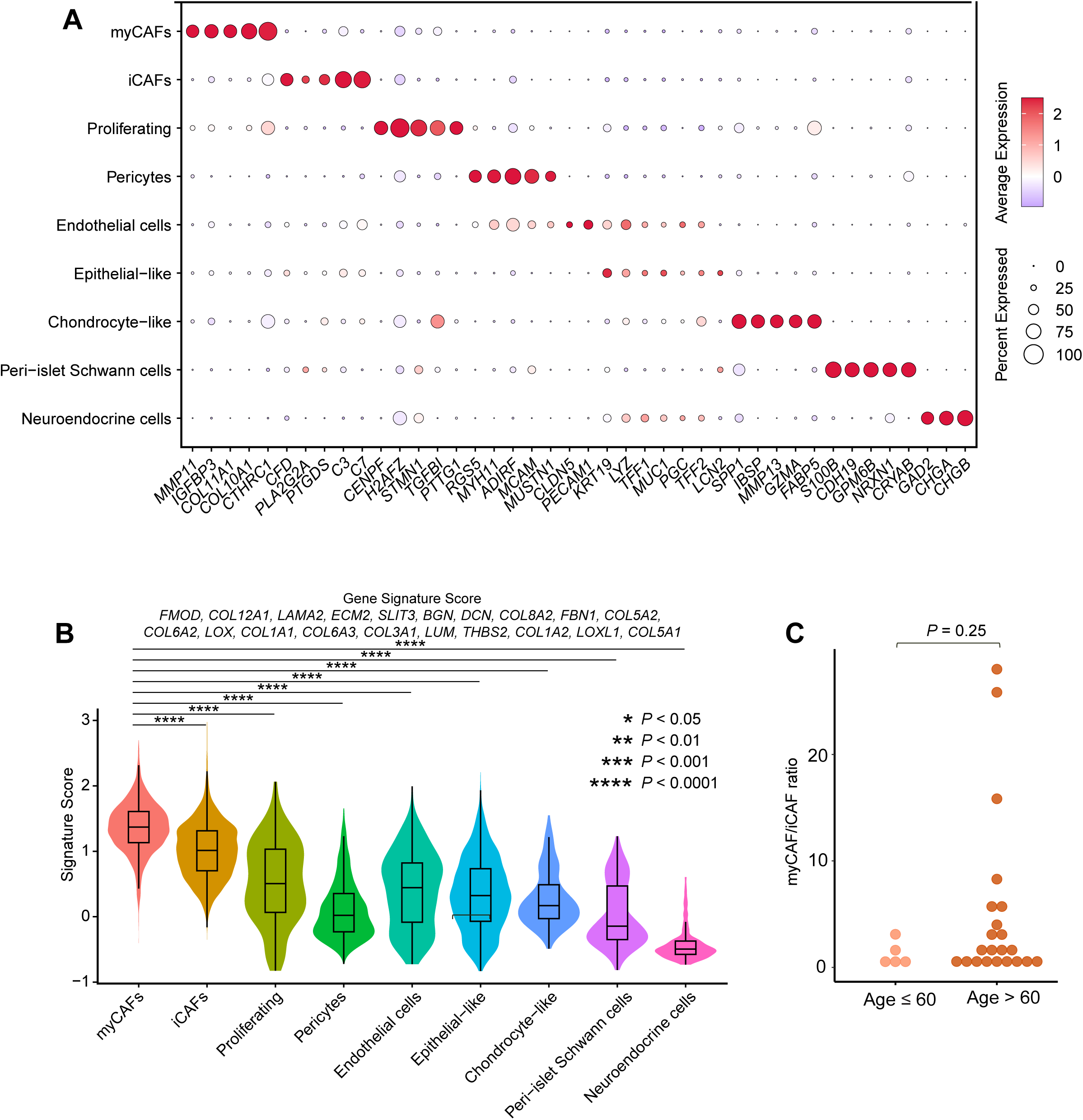
Charting ECM gene signature across CAF subtypes in human PDAC. **A**, Dotplot showing scaled expression levels of marker genes used to identify CAF subtypes obtained from re-clustering of CAF cells in human PDAC datasets from *Werba et al.* **B**, Signature score of our 20-gene ECM signature in the CAF subtypes. Significance levels (Wilcoxon rank sum test) for myCAF versus other subtypes are indicated. **C**, myCAF/iCAF cell ratios for old and young patients are indicated.

**Supplementary Figure 5.**
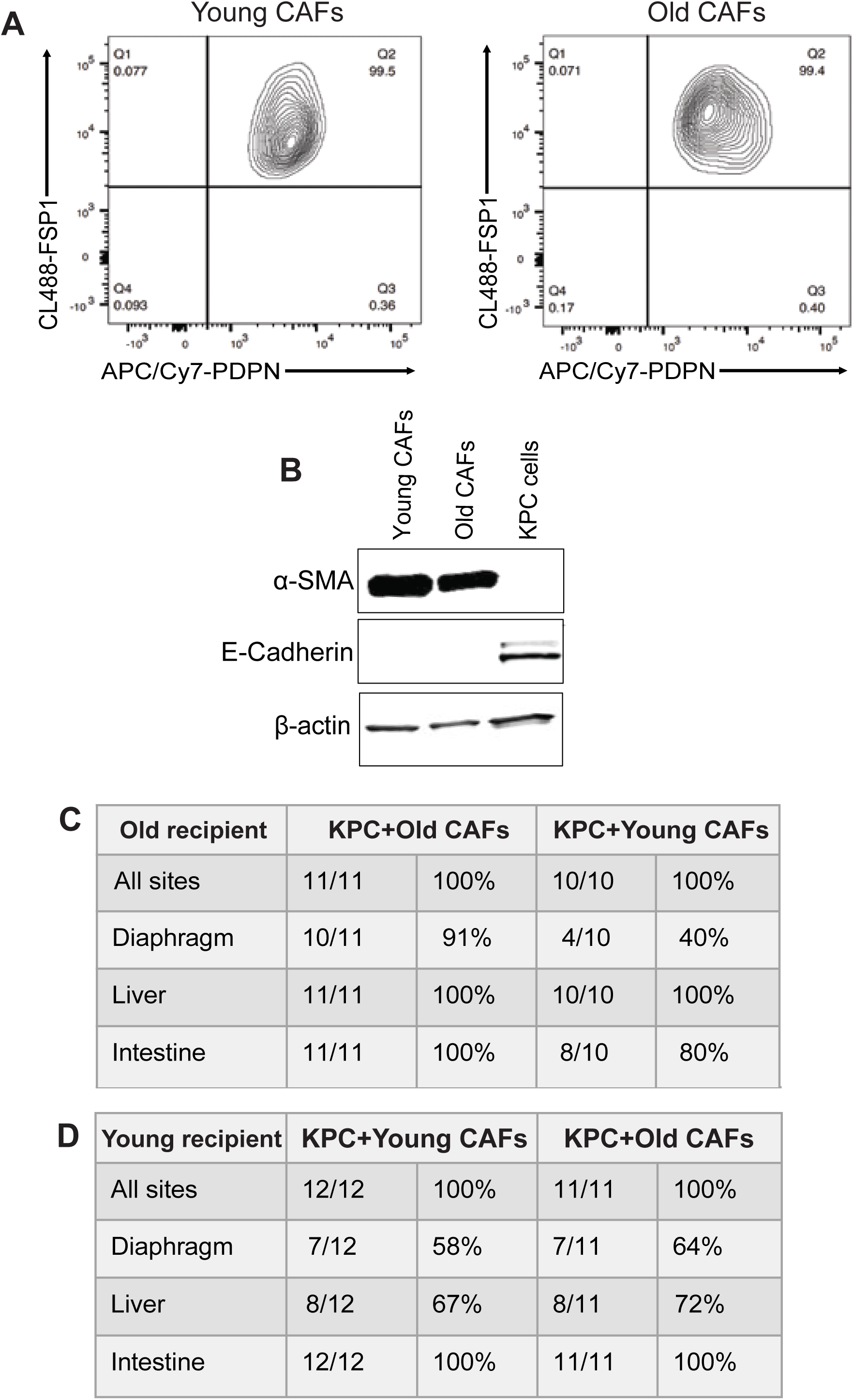
Heterochronic restructuring of the tumor stroma alters age-dependent metastatic effects. **A,** Flow cytometry-based characterization of isolated murine CAFs for pan-CAF markers, including podoplanin (PDPN) and fibroblast-specific protein 1 (FSP1). **B,** Immunoblot showing the expression of fibroblast marker α-SMA in young and old CAFs, and the epithelial marker E-Cadherin in KPC cells. β-actin serves as a loading control. **C−D**, Table summarizes the incidence of metastases across various organ sites in old and young recipients, following heterochronic co-implantation of murine CAFs with KPC cells.

